# Heterochromatin drives organization of conventional and inverted nuclei

**DOI:** 10.1101/244038

**Authors:** Martin Falk, Yana Feodorova, Natasha Naumova, Maxim Imakaev, Bryan R. Lajoie, Heinrich Leonhardt, Boris Joffe, Job Dekker, Geoffrey Fudenberg, Irina Solovei, Leonid Mirny

## Abstract

The mammalian cell nucleus displays a remarkable spatial segregation of active euchromatic from inactive heterochromatic genomic regions. In conventional nuclei, euchromatin is localized in the nuclear interior and heterochromatin at the nuclear periphery. In contrast, rod photoreceptors in nocturnal mammals have inverted nuclei, with a dense heterochromatic core and a thin euchromatic outer shell. This inverted architecture likely converts rod nuclei into microlenses to facilitate nocturnal vision, and may relate to the absence of particular proteins that tether heterochromatin to the lamina. However, both the mechanism of inversion and the role of interactions between different types of chromatin and the lamina in nuclear organization remain unknown. To elucidate this mechanism we performed Hi-C and microscopy on cells with inverted nuclei and their conventional counterparts. Strikingly, despite the inversion evident in microscopy, both types of nuclei display similar Hi-C maps. To resolve this paradox we developed a polymer model of chromosomes and found a universal mechanism that reconciles Hi-C and microscopy for both inverted and conventional nuclei. Based solely on attraction between heterochromatic regions, this mechanism is sufficient to drive phase separation of euchromatin and heterochromatin and faithfully reproduces the 3D organization of inverted nuclei. When interactions between heterochromatin and the lamina are added, the same model recreates the conventional nuclear organization. To further test our models, we eliminated lamina interactions in models of conventional nuclei and found that this triggers a spontaneous process of inversion that qualitatively reproduces the pathway of morphological changes during nuclear inversion *in vivo*. Together, our experiments and modeling suggest that interactions among heterochromatic regions are central to phase separation of the active and inactive genome in inverted and conventional nuclei, while interactions with the lamina are essential for building the conventional architecture from these segregated phases. Ultimately our data suggest that an inverted organization constitutes the default state of nuclear architecture.

---

Mammalian interphase chromosomes are spatially organized with distinct features ranging from chromosome territories, to active and inactive A/B compartments and to TADs^1,2^. Emerging work argues that these features reflect different underlying mechanisms^3^. Of much recent interest^4–8^, TAD formation appears to result from a cohesin-mediated mechanism of loop extrusion. Compartments are increasingly appreciated as distinct from TADs: compartments both persist when TADs are experimentally disrupted^5,6,8,9^, and can be absent when TADs are present (e.g. in mouse maternal zygotes^10^). Still, the mechanisms that underlie the segregation of active and inactive chromatin, leading to spatial nuclear compartmentalization, remain unknown.

In conventional mammalian nuclei, microscopy and chromosome conformation capture techniques provide complementary insight into the spatial segregation and compartmentalization of active and inactive regions. Microscopy reveals that transcriptional state is tightly related to spatial compartmentalization: inactive heterochromatic loci are found either at the nuclear periphery, associated with the nuclear lamina and chromocenters, or adjacent to nucleoli; active euchromatic loci, on the contrary, are found in the nuclear interior^11,12^. Hi-C reveals that this segregation occurs genome-wide, and manifests as a plaid pattern of enriched contact frequency^2^. This plaid pattern reflects (i) presence of at least two types of chromosomal regions, termed A- and B-compartments, and (ii) about two-fold enrichment of contacts between regions that belong to the same compartment type, which is evident both within (cis) and between (trans) chromosomes. Consistent with imaging, A-compartment regions identified by Hi-C carry euchromatic histone marks, are gene rich, early-replicating and transcriptionally active; B-compartments are enriched in heterochromatic marks, gene poor, late-replicating, and frequently associated with the nuclear lamina^12^. These close associations have led to numerous proposals for causal roles of the lamina^12^, and mobility^13–15^- or transcription^16,17^-related clustering of euchromatin in mediating A/B compartmentalization. Still, despite these insights, which type of chromatin or what kind of nuclear structures causally underlie compartmental segregation remains unknown.

The inverted nuclei of nocturnal mammalian rods, which exhibit a dramatic departure from this conventional pattern of compartmentalization, provide an opportunity to elucidate mechanisms underlying spatial compartmentalization. Specifically, inverted rod nuclei in mouse have a strikingly regular concentric organization: pericentric constitutive heterochromatin (major satellite repeat) is packed into a single central chromocenter, which is encircled by a shell of facultative heterochromatin (LINEs/LTRs-rich) and then by an outermost shell of euchromatin (Fig. 1a)^18^. Constitutive and facultative types of heterochromatin are functionally different as they carry different combinations of epigenetic marks, which remain preserved through the inversion^19^, suggesting that only spatial locations rather than composition of different chromatin types change upon inversion. We hypothesized that this natural perturbation could provide a rich testing ground for mechanistic theories of compartmentalization.

**Figure 1.**
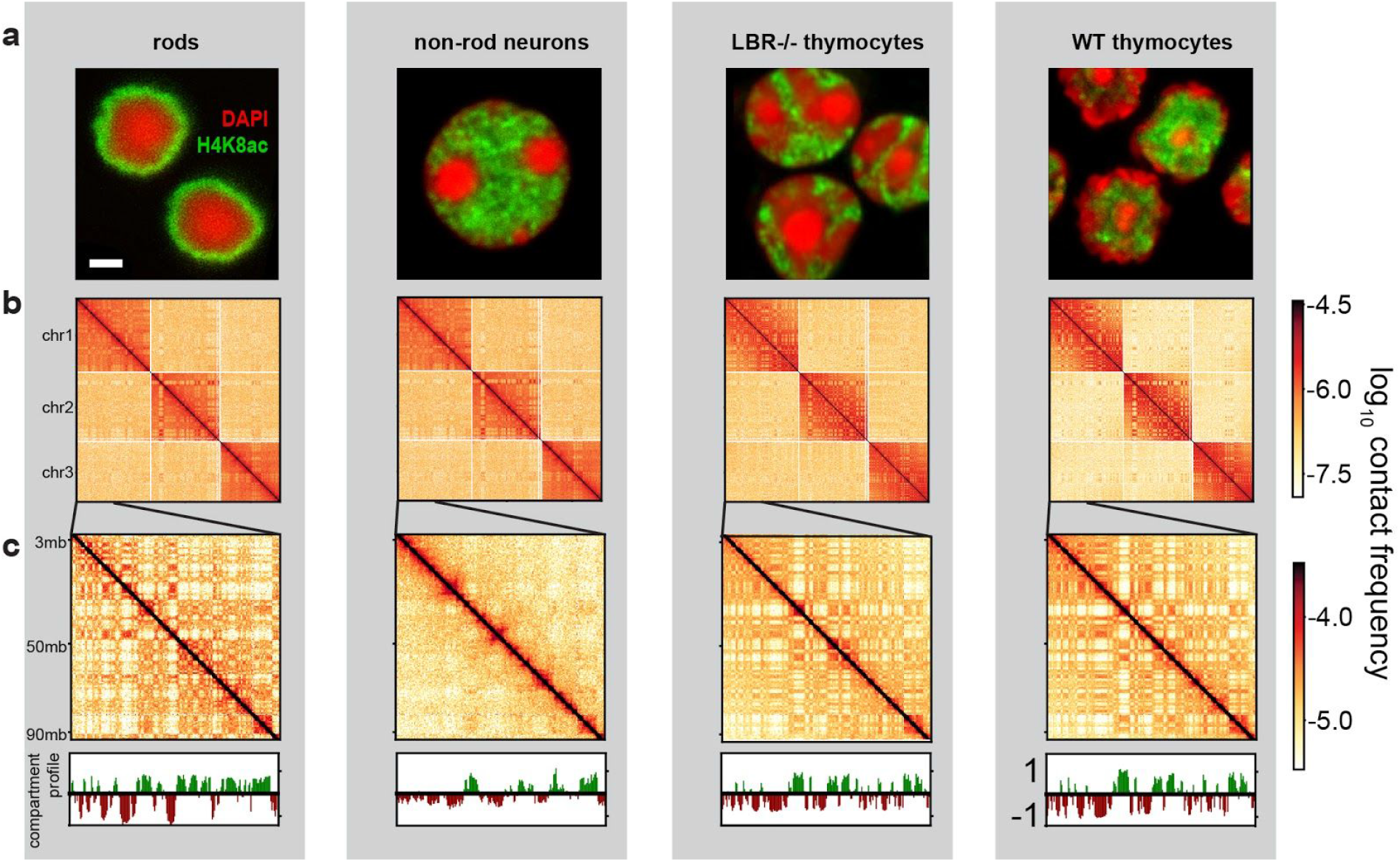
Nuclear architecture of the studied cell types revealed by microscopy and Hi-C. **a**, The spatial organization of eu- and heterochromatin are revealed by staining with anti-H4K8ac antibody (green) and DAPI (red). Nuclei of non-rod neurons and WT thymocytes are conventional, with euchromatin in the interior. Rod nuclei are inverted, with a single central chromocenter. Nuclei of LBR-null thymocytes are inverted and have several chromocenters. Scale bar, 2 μm. **b**, Hi-C contact maps for chromosomes 1, 2, and 3 show a checkerboard pattern in cis and trans, reflecting compartmentalization, with cis contacts much more frequent than trans contacts. **c**, Hi-C contact maps for an 87MB region of chr1, with compartment profile indicating regions in the A (green) and B (brown) compartment above.

Previous models of compartmentalization have suggested several possible mechanisms, including preferential attraction of similar chromatin to each other, such as A-A and B-B attraction^3,20–23^, attraction of heterochromatin to the nuclear lamina^23^, and higher level of chromatin mobility in the active chromatin^13–15^. Still, mechanistic proposals regarding how compartments are formed and positioned remain inconclusive due to the difficulty of disentangling possible contributing factors within conventional nuclei^24^.

To understand general mechanisms of genome compartmentalization we performed Hi-C in four mouse cell types isolated from primary tissues that have either conventional or inverted nuclear architectures: rod photoreceptors (inverted), non-rod retinal neurons (conventional), WT thymocytes (conventional), and LBR-null thymocytes^19,25^ (inverted) (Fig. 1a; Table S1). The latter three cell types provide excellent points of comparison to rods: retinal non-rod neurons are similarly post-mitotic but have large conventional nuclei; thymocytes are cycling cells with nuclei of a similar size.

Surprisingly, despite the great differences in nuclear organization evident from microscopy (Fig. 1a), all features of chromatin organization characteristic of conventional nuclei - TADs, chromosome territories, and compartments – are present in inverted nuclei, although with quantitative differences (Fig. 1b,c; Extended Data Fig. 1). However, these difference stem from differences between cell types and do not appear to be related to nuclear inversion. TAD strength is the lowest in rods, highest in non-rod neurons, and intermediate in both WT and LBR-null thymocytes (Fig. 2a,c), as measured using annotated TADs from murine ESCs^26^ (Methods) and by computing insulation profiles (Extended Data Fig. 1). The comparable TAD strengths and insulation profiles of LBR-null and WT thymocytes suggests that TADs are unaffected by inversion, and furthermore that the difference in TAD strength of non-rod neurons and rods must be due to other aspects of nuclear physiology, unrelated to inversion. Chromosome territoriality is a large-scale feature of a Hi-C map, manifesting itself as an increased fraction of within-chromosome (cis) contacts (Methods). Territoriality is stronger in WT and LBR-null thymocytes and weaker in non-rod and rod neurons (Fig. 2e), consistent with the more dispersed chromosomes observed microscopically in both rod and non-rod neurons (Fig. 2f), and is thus not a function of the nuclear inversion.

**Figure 2.**
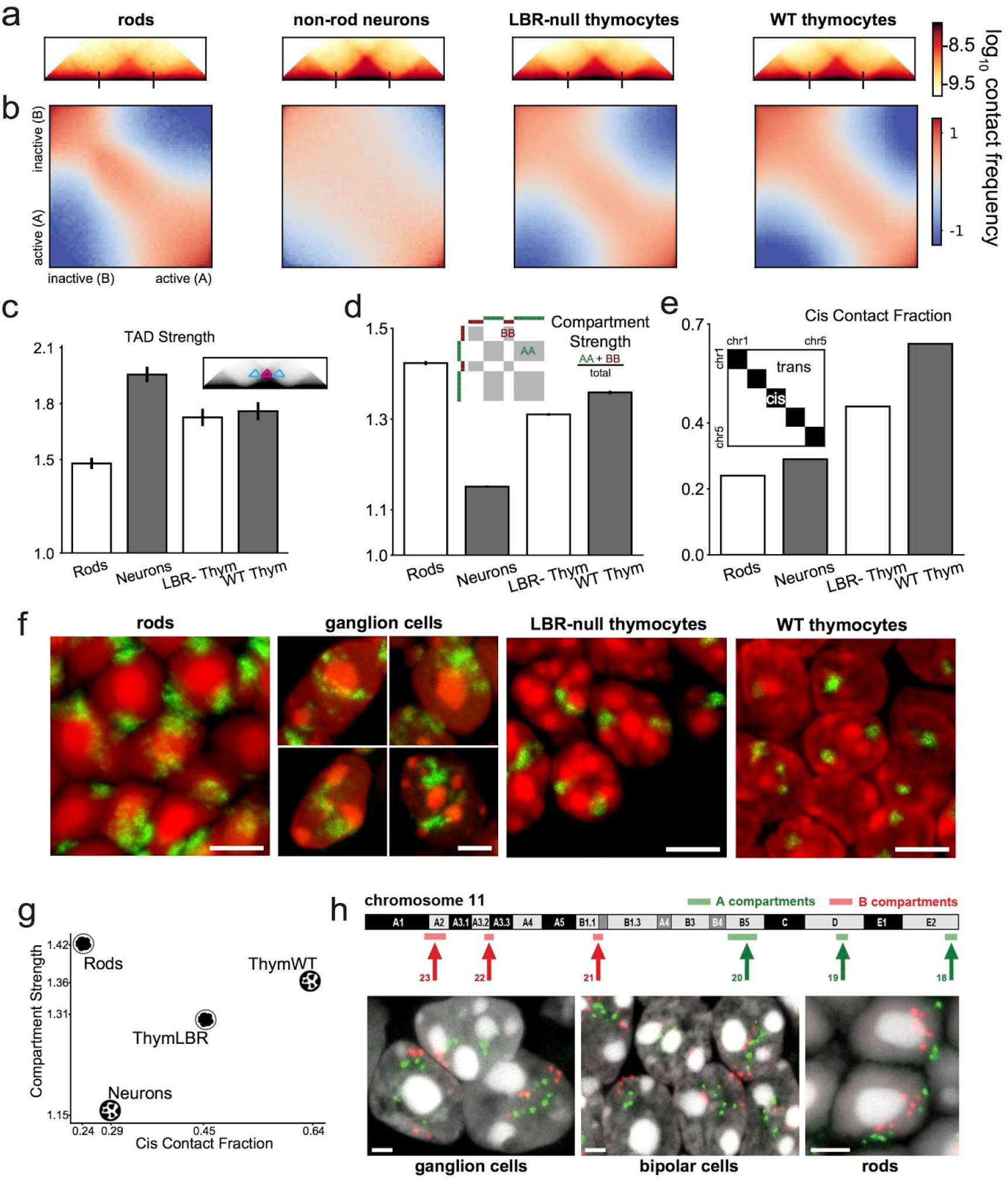
Quantification and microscopic verification of the strength of Hi-C features. **a**, Average TADs, based on domain calls from ESC^5^. Ticks indicate start and end of TADs. **b**, Saddle plots^27^ (see Methods) displaying the extent of compartmentalization across cell types. **c**, TAD strength is weakest in rods and strongest in non-rod neurons. TAD strength is the ratio of average contacts within the TAD (blue triangle on the inset) to average contacts between TADs (pink triangles). **d**, Compartmentalization is strongest in rods and weakest in non-rod neurons, with the schematic indicating the two regions of contacts compared to quantify compartmentalization. **e**, Chromosome territoriality, measured as the ratio of *cis* contacts to *cis+tans* contacts, is weaker in rods and non-rod neurons in comparison to conventional and inverted thymocytes. The schematic illustrates the compared regions. **f**, Consistent with the low *cis* contact fraction revealed by Hi-C, chromosome 11 visualized by FISH (green) has a more diffuse territory in postmitotic rods and non-rod neurons in comparison to cycling thymocytes of both genotypes. Projections of 2 μm confocal stacks; scale bars, 5 μm. **g**, Scatterplot of compartmentalization and territoriality shows these features are not directly related. **h**, Inverted localization of A and B compartment loci in rods compared to non-rod neurons. FISH with six BACs hybridizing to constitutive A (green) or B (red) compartments of chromosome 11 (for BAC coordinates see Methods). Projections of 3 μm confocal stacks; scale bars, 2 μm.

Most surprisingly, the lack of lamina association in inverted nuclei does not compromise their compartmentalization. This refutes the proposal (for review^12^) that tethering of heterochromatin to the lamina underlies compartmentalization. We computed compartment profiles from Hi-C maps using eigenvector decomposition^27^ and defined the degree of compartmentalization as the enrichment of contacts between compartments of the same type (Methods). While assignments of A/B-compartments are generally cell-type dependent^2^, compartment profiles remain highly correlated upon perturbing lamina association in thymocytes *(r(LBR,WT)=.95; p<10^−5^*, Extended Data Fig. 1). Moreover, the degree of compartmentalization remains unchanged in thymocytes upon inversion, and even becomes stronger in rods with a higher degree of inversion. Taken together, our analyses show that major features of nuclear organization, in particular the degree of compartmentalization, are qualitatively preserved despite the nuclear inversion.

It is also striking that Hi-C maps are not informative of the global 3D reorganization of the nucleus upon inversion, which is obvious in microscopy (Fig. 2d,h). By measuring contact frequencies rather than distances or spatial locations, Hi-C provides information largely complementary to that obtained with microscopy^28^. This calls for the development of a model of nuclear compartmentalization that synthesizes the orthogonal data from Hi-C and imaging. Moreover, the lack of functional chromatin-lamina tethers (both LBR- and LamA/C-dependent) provides an important opportunity to disentangle the role of the lamina from that of interactions between chromatin regions in establishing nuclear organization, and to seek for universal mechanisms that establishes compartmentalization in both conventional and inverted nuclei.

We sought a mechanism of compartmentalization that is able to recapitulate (i) the inverted organization in the absence of interactions with the lamina, reproducing both the architectures we observe microscopically and the compartment strength measured from Hi-C data, and (ii) the conventional architecture and the Hi-C compartment strength by adding attractive interactions between heterochromatin and the nuclear lamina. As small-scale features like TADs were unrelated to inversion (Fig. 2c), we focused on developing models sufficiently coarse-grained to recapitulate whole-scale nuclear geometry. Following this mechanistic *de-novo* approach^29^, we first developed a model for the inverted nucleus.

Similarly to other phase-separation models of compartmentalization^20–23^, we represented chromosomes as block copolymers with three types of monomers, and studied their equilibrium organization (Fig. 3a). In our simulations, each monomer can be of one of three types - euchromatin (A), heterochromatin (B) or pericentromeric constitutive heterochromatin (C) - and corresponds to ~30 Kb of DNA. We generated a polymer representation of 8 chromosomes, each consisting of 6000 monomers (180 Mb total), confined to a spherical nucleus (density 35%). Monomers have excluded volume cores, and can attract each other depending on their chromatin type. The sequence of A and B monomers along the polymer mirrors the sequence of A and B compartments derived from Hi-C data of rods (Fig. 3a; Methods). To simulate the acrocentric structure of mouse chromosomes, we placed a block of C monomers constituting 10% of each chromosome at one end of each polymer. We consider B and C heterochromatin separately as they are distinct in microscopy; pericentric constitutive heterochromatin consists mostly of satellite repeat sequences, forms chromocenters evident in microscopy, but is unmappable via Hi-C. For each parameter set considered below, we generated an equilibrium ensembles of polymer conformations, and analyzed their nuclear geometries and compartmentalization to compare with those obtained from experimental microscopy and Hi-C.

**Figure 3.**
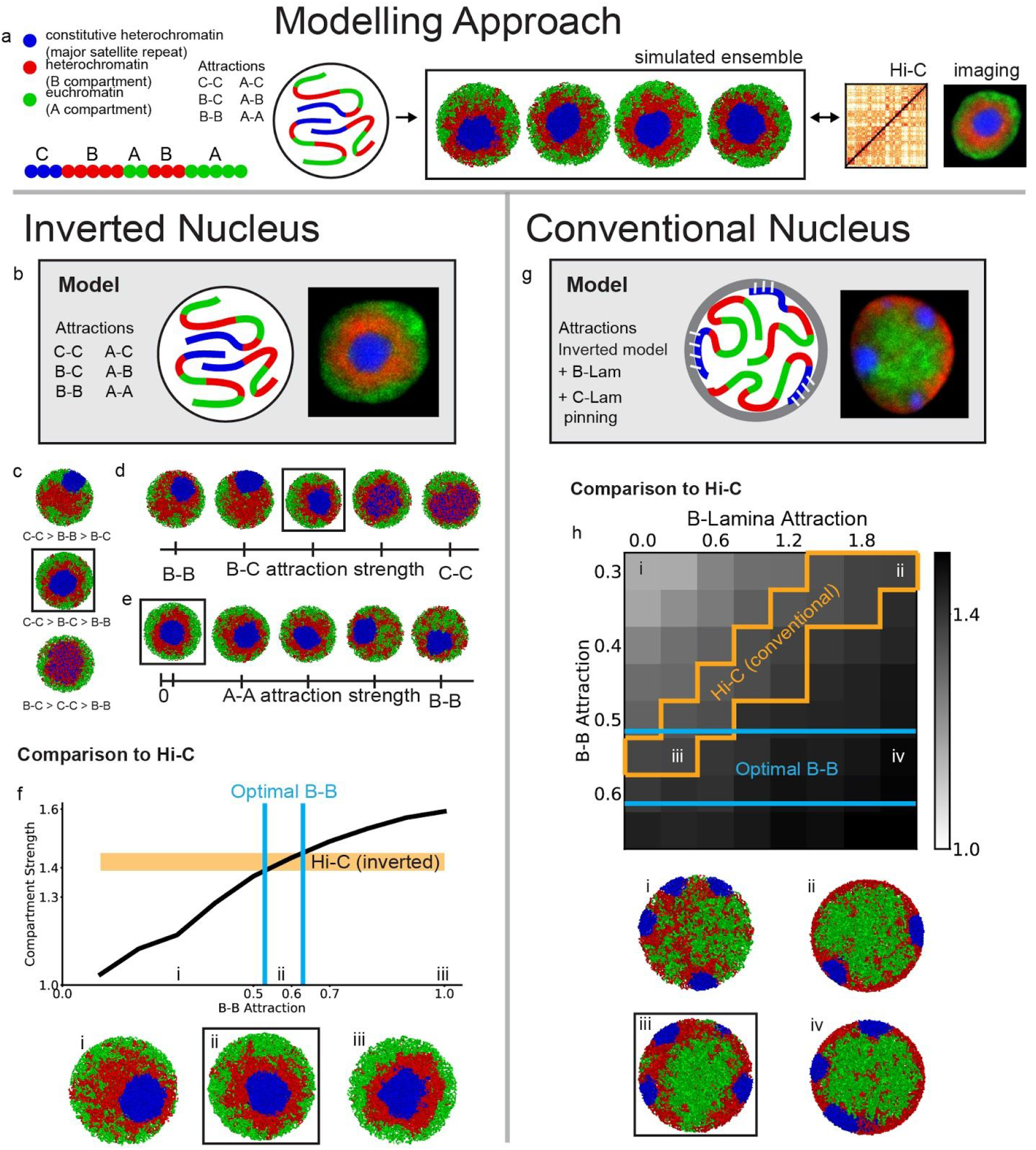
Polymer model reproduces microscopic morphologies and strength of Hi-C features. **a**, Our approach is to: define a mechanistic model with parameters describing chromatin interactions; simulate an ensemble of configurations predicted for this model via Langevin dynamics; and compare these configurations to Hi-C and microscopy data. In the panels below, we show a representative conformation for the indicated parameter set; for calculating compartment strength, we compute the ensemble average. **b-f, model of the inverted nucleus**. **b**, Each chromosome is represented as a heteropolymer, consisting of C (blue), B (red), and A (green) monomers, corresponding to constitutive heterochromatin, heterochromatin and euchromatin, respectively. The model parameters are based on the attraction between every pair of monomer classes. A representative microscopic image of a rod nucleus is shown on the right. **c**, The three attraction strengths, C-C, B-C and B-B, can be ordered only in one way (middle configuration) to faithfully reproduce the morphology of the rod nucleus. **d**, Varying the value of B-C attraction between that of B-B and C-C does not affect the inverted nuclear morphology, up until a certain value at which B-C is roughly (B-B+C-C)/2. This value is what we should expect based on naïve application of the Flory-Huggins mixing theory. We choose to set B-C value to be the geometric mean of B-B and C-C, below this threshold (boxed configuration). **e**, Varying A-A interactions from 0 to that of B-B produces different morphologies in terms of the visible separation of A and B monomers, but not in any way that cannot be replicated simply by keeping A-A to 0 and tuning the value of B-B. We choose to set A-A at a value slightly above zero and much less than B-B (boxed configuration). **f**, Varying the B-B attraction in simulations of inverted nuclei identifies a range for which experimental compartment strength can be matched. Particular values of B-B are selected and snapshots of simulations at those values are shown. The orange bar indicates experimental range of compartment strength in rods between two replicate experiments of Hi-C. Vertical lines indicate values of B-B attraction where simulated compartment strength passes through the experimental range. **g-h model of the conventional nucleus**. **g**, The model for conventional nuclei additionally includes interactions of monomers with the nuclear lamina. B monomers are attracted to the lamina with a strength B-Lam and C monomer clusters are pinned to the lamina at random positions. A representative microscopic image of a conventional nucleus is shown on the right. **h**, Compartment strength as a function of B-B and B-Lam attractions. Particular points in this two-dimensional parameter space are selected and snapshots of those simulations are shown below. Orange lines indicates regions in parameter space where simulated Hi-C has compartmentalization close to (within .03 of) that found in experimental Hi-C. Note that experimental compartment strength can be matched even if we constrain B-B interactions to be in the same range as for inverted nuclei (at point iii).

We started by finding a class of models that can reproduce the geometry of the inverted nucleus. While our model has six parameters that represent the attraction between every pair of monomer classes (Fig. 3b), the need to reproduce the inverted architecture with concentric regions of C, B, and A chromatin reveals crucial and strong constraints on the ordering of the attraction energies (Fig. 3c-e). First, recapitulation of the single central chromocenter required setting the C-C attraction as the strongest. Models where B and C monomers were set to be identical, effectively a two-type model, were unable to reproduce a separate C core surrounded by a B layer. Second, the surrounding shell of heterochromatin necessitated B-B to be weaker than B-C, otherwise, B-heterochromatin expelled the chromocenter to the nuclear periphery, minimizing the B-C interface. Similarly, to reproduce the outer shell of euchromatin, A-A attraction had to be weaker than A-B. Moreover, having clearly resolved and narrow interfaces between different types of chromatin (e.g. A-B and B-C interfaces) required attraction between different chromatin types to be in between homotypic attraction strengths (i.e. A-A<A-B<B-B; B-B<B-C<C-C), consistent with similar histone marks in B and C regions. As such, the A-B attraction was fixed to be the geometric mean of A-A and B-B, B-C to be that of B-B and C-C, and A-C to be that of A-A and C-C. Furthermore, we do not find alternate morphologies in simulations with small A-A attraction strengths (Fig. 3e), and hence subsequently fix A-A a value much smaller than BB. Overall, simulations that could reproduce an inverted geometry required a particular order of interaction strengths A-A<A-B<B-B<B-C<C-C (Fig. 3c-e), with B-B attraction energy as the only free parameter in our model.

To simultaneously reproduce the compartmentalization measured in Hi-C data and the inverted organization seen in microscopy, we swept across possible values of B-B attraction energy. We found a narrow range (0.5–0.6kT) for which our model is quantitatively consistent with both Hi-C and microscopy in rods (Fig. 3f). Thus, our models indicate that attraction between heterochromatic regions can generate compartmentalization in agreement with Hi-C and global architecture that of inverted nuclei as seen in microscopy. The central role for attractions between heterochromatic regions is in contrast to suggestions from earlier studies concentrated on the role of the lamina^12^ and activity-related clustering of euchromatin^13–17^. Importantly, we found that interactions between constitutive heterochromatin (*C*, H3K9m3-rich) should be considerably stronger than interactions between facultative heterochromatin (*B*, H3K27me3/H4K20me3-rich). As a number of histone marks are shared between constitutive (C) and facultative (B) heterochromatin^19^, these marks may directly control chromatin interaction strength, with B heterochromatin simply having lower quantitative levels of these marks, consistent with the observation of marks that are present in B, yet absent in C. Our model is agnostic of whether such attraction between heterochromatin is caused by affinity between homotypic repetitive elements^30,31^, or mediated by other proteins (e.g. HP1)^32,33^. Recent observations on heterochromatin demixing from the entire nuclear chromatin by phase separation^32,33^ additionally support the essential role of heterochromatic interactions in chromatin compartmentalization. Equally importantly, our model shows that interactions within euchromatin are nonessential for compartmentalization.

To extend our model to conventional nuclei, we incorporated an attraction of heterochromatin to the nuclear lamina (Fig. 3g), consistent with the role of lamina-associated domains in nuclear organization^12^. While B-monomers are attracted to the lamina with the energy Lam-B, C monomers were tethered to the nuclear lamina by stronger (irreversible) bonds (Fig. 3g), modelling the lamina-associated distribution of chromocenters. By varying both B-B and B-Lam attractions, we found that our model is capable of reproducing both radial ordering of active and inactive chromatin from microscopy and compartmentalization from Hi-C data in WT thymocytes for a range of B-B and B-Lam values. Importantly, histone modifications remain associated with the same class of chromatin in both inverted and conventional nuclei^19,25^. Based on this biological evidence, we parsimoniously assume that B-B attraction remains the same in conventional and inverted nuclei. With this constraint, we can narrow the range of possible B-Lam values (~0.5kT) (Fig. 3h) and find that B-Lam attraction should be comparable to B-B attraction. We note that our models can be readily extended to account for subtleties of locus-specific chromosome compartmentalization that future experiments may uncover, e.g. including potentially differential behaviors of fLADs, cLADs^12^, and NADs^34^, cell-to-cell variation in LAD presence at the lamina ^35^, and interactions of chromatin with other nuclear bodies (polycomb^36^, speckles^37^, etc.). Together, our results support the essential role of peripheral heterochromatin tethers in establishing conventional nuclear architecture^12,25,38^.

We then tested our model by comparing its predictions with a microscopy time-course through the different stages of nuclear inversion. Specifically, we tested whether elimination of peripheral heterochromatin tethers is sufficient for the induction of nuclear inversion^25^. In agreement with this, when we began simulations in a conventional organization and then turned off lamina-heterochromatin interactions, the simulated nuclei slowly became inverted (Fig. 4a,b). While we expect the default, inverted state to emerge upon disruption of the lamina, there is no guarantee that the simulated pathway by which this transformation takes place is similar to that seen in real neurons. Remarkably, the changing geometry of simulated nuclei closely mirrored the changes seen during rod differentiation^18,25^, with B and C monomer droplets undergoing a liquid-like fusion, characteristic of a phase separation and similar to *in vitro* phase-separated systems^32,33,39^ (Fig. 4a,b). Although compartmentalization dips after heterochromatin moves away from the lamina (Fig. 4a), compartments remain separated during the whole process of inversion. To confirm this prediction, we performed additional microscopy, and found that small chromosome segments move together with chromatin of their own compartment type during nuclear inversion (Fig. 4c, Extended Data Fig. 2). In particular, the rhodopsin gene undergoes long-range movements from the interior to the periphery of the nucleus during rod differentiation, but remains within the active euchromatic compartment (Fig. 4d1). Correspondingly, synthesis of rhodopsin, which starts in the still conventional nuclei of rod progenitors, continues and increases concomitantly with nuclear inversion (Fig. 4d2). Thus, the dramatic reorganization of nuclear architecture accompanying rod differentiation does not impede segregation or function of eu- and heterochromatin^11^.

**Figure 4.**
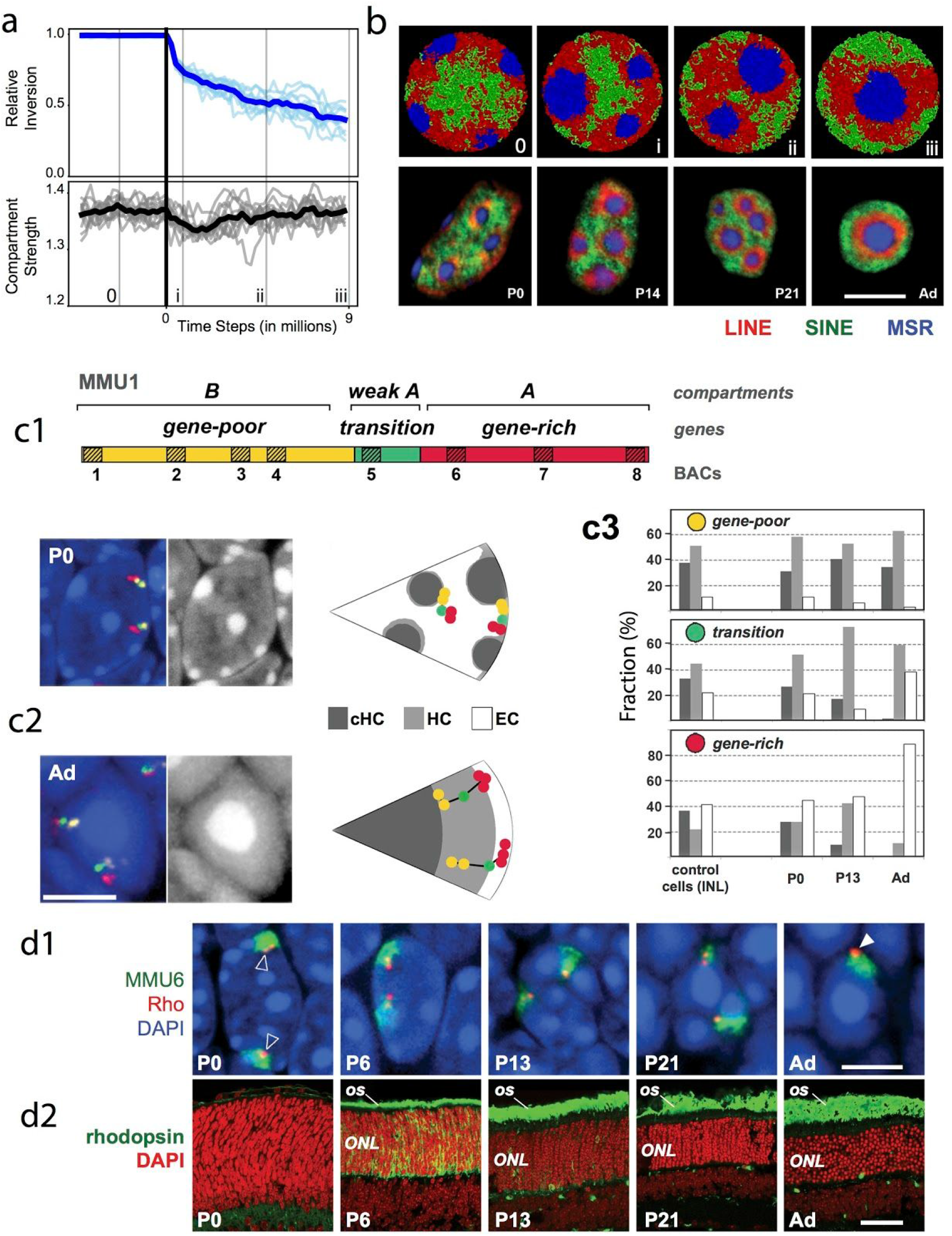
Nuclear inversion does not disrupt compartmentalization. **a**, Simulated time-course of the process of nuclear inversion. Configurations from particular time points indicated by numerals and thin lines are displayed in (b). Solid vertical line indicates the time at which interactions with the lamina are eliminated. (top) Relative to their initial positions, C monomers move towards the nuclear interior following lamina interaction removal. Light blue lines are computed from individual simulations, the blue line is the average of these simulations. (bottom) There is only a small dip in compartmentalization of simulated Hi-C following lamina interaction removal. Grey lines are computed from individual simulations, the black line is the average of these simulations. **b**, Simulations (*top raw*) faithfully reproduce the change in chromatin architecture that take place during rod differentiation *in vivo* (*bottom raw*). Single confocal sections after FISH with probes for LINE1 (red), B1 (green) and major satellite (blue). *P0-P21*, postnatal days; *Ad*, adult. Scale bar, 5 μm. **c**, Validation of the maintenance of compartmentalization during inversion by microscopy. Chromosomal loci follow the spatial position of the compartment they are associated with during differentiation. **c1**, Schematics of the subchromosomal regions belonging to either A (red) or B (yellow) compartments of chromosome 1. The regions are labeled by BAC probes (1–8) depicted as striped boxes. **c2**, Representative images of rod nuclei in P0 and adult retina after 3-color FISH and accompanying schemes showing typical distribution of BAC signals. 2 μm projections of confocal stacks. Scale bar, 5 μm. **c3**, Analysis of A and B loci distribution in the different compartments - constitutive heterochromatin of chromocenters (dark grey), heterochromatin (light grey) and euchromatin (white) - at three stages of rod differentiation and in control cells (non-rod neurons). Note relocation of A loci from the nuclear interior to the nuclear periphery during rod maturation. Histograms show proportions of loci signals in three compartments. For other example and detailed description of the experiment, see Extended Data Fig. 2). **d**, Repositioning of genomic loci together with their compartment coincides with maintenance of transcriptional activity. **d1**, In the process of nuclear inversion, rhodopsin locus (red) within chromosome 6 (green) changes position from internal (empty arrowheads) to peripheral (solid arrowhead). **d2**, Despite this dramatic movement, rhodopsin expression (green) in rods, which starts in still conventional rod nuclei, continues at an increasing rate concomitantly with the nuclear inversion. 2 μm projections of confocal stacks; scale bars, 5 μm (d1) and 50 μm (d2). BAC probes for the rhodopsin locus are listed in Supplementary Table S1. *OS*, outer segments of rods positive for rhodopsin staining; *ONL*, outer nuclear layer containing rod perikarya.

Together, our results show the central role of interactions between heterochromatin in establishing compartmentalization by phase separation, and the role of tethering to the lamina in spatial locations of compartments in conventional nuclei. We found that Hi-C maps for naturally and experimentally inverted nuclei remarkably resemble those of conventional nuclei, effectively capturing their similarly compartmentalized topology, despite strikingly different geometry of their 3D organization. Taking advantage of complementary microscopy and Hi-C data, we developed an integrated model for the two contrasting nuclear architectures, which shows that (i) interactions between heterochromatin regions lead to phase separation of chromatin and are essential for the compartmentalization of conventional and inverted nuclei, (ii) lamina-heterochromatin interactions are dispensable for segregation of eu- and heterochromatin, but central in establishing the conventional nuclear architecture, and (iii) euchromatic interactions are dispensable for compartmentalization. This mechanism reconciles two complementary views of nuclei as seen by Hi-C and microscopy. Taken together, our results indicate that the inverted nucleus, with its phase-separated heterochromatic center, constitutes the default nuclear architecture imposed by the mechanism of compartmental interactions and must be complemented by lamina-heterochromatin interactions to form the conventional nucleus. Since inverted nuclei in rods and lymphocytes are fully functional, our work raises the questions of why most eukaryotic nuclei have conventional organizations, and what is the role of the peripheral location of heterochromatin.

## Data availability

Hi-C maps can be browsed and compared in HiGlass^40^ browser at http://mirnylab.mit.edu/proiects/invnuclei/ and at a public server http://higlass.io/app/?config=JLOhiPILTmq6qDRicHMJqg

## Acknowledgments

We are grateful to all member of the Mirny lab for many useful discussions, and to Nezar Abdennur and Peter Kerpedjiev for help with HiGlass Hi-C browser. This work has been supported by NSF 1504942; NIH GM114190; NIH HG003143, and NIH DK107980 grants. MF was supported by the Department of Defense (DoD) through the National Defense Science & Engineering Graduate Fellowship (NDSEG) Program; by Deutsche Forschungsgemeinschaft to IS (SO1054/3 and SFB1064).

